# Copper Stress Trigger Organelles Communication and Chromatin Condensation Leading to Cell Death in *Solanum lycopersicum*

**DOI:** 10.1101/2025.07.17.665307

**Authors:** Sakshi Chouhan, Shilpa Chandra, Abdul Salam, Chayan Kanti Nandi

## Abstract

Copper (Cu) is a vital micronutrient for plants but becomes highly toxic when present in excess, disrupting redox balance and damaging cellular structures. While the physiological and mitochondrial responses to Cu toxicity are well-documented, the nuclear-level consequences, particularly chromatin remodeling and gene regulatory changes, remain poorly understood. In this study, we used the root apex of *Solanum lycopersicum* as a model system to explore how increasing copper concentrations affect organelle integrity, stress signaling, and nuclear architecture. Using confocal and super-resolution imaging with organelle-specific markers and immunostaining, we observed that mitochondria was the earliest affected to Cu stress, exhibiting fragmentation, membrane depolarization, and cytochrome c release led to reactive oxygen species (ROS) accumulation, activation of AMPK, suppression of mTOR signaling, and nuclear translocation of NRF2. Critically, we found that copper exposure induced profound nuclear alterations, including shrinkage, lobulation, peripheral chromatin tethering, and global condensation events tightly correlated with stress signaling. H3K4me3 immunostaining revealed a shift from active euchromatin to condensed, transcriptionally silent states, leading to membrane rupture and cell death in root tip cells. Our findings show that the nucleus actively integrates organelle-derived stress signals, with chromatin remodeling as a key marker of copper toxicity. This highlights potential for chromatin-based diagnostics and stress-resilient crop breeding.

**Graphical Abstract:** 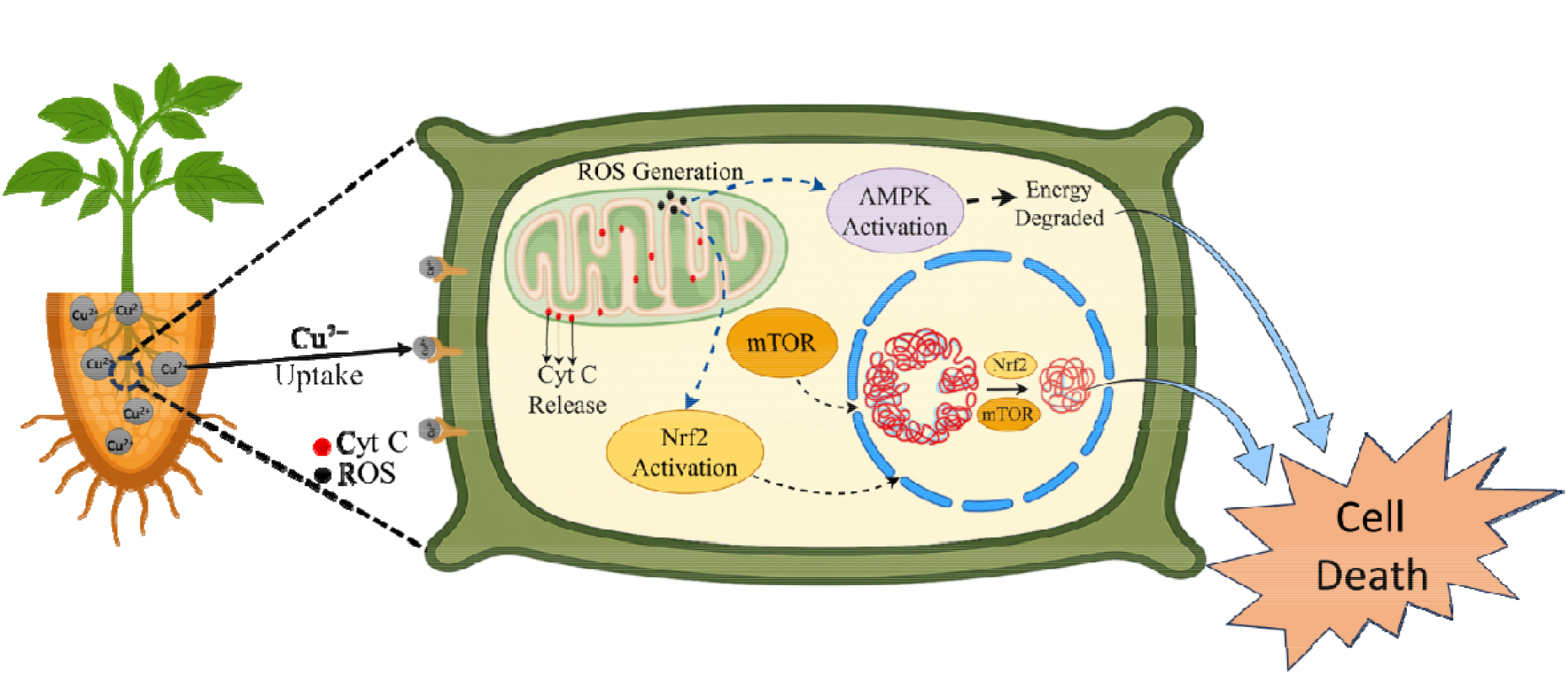

## Introduction

Copper (Cu) is an essential micronutrient required for numerous physiological and biochemical processes in plants, including mitochondrial respiration, antioxidant defence, iron mobilization, and photosynthetic electron transport ^1–5^. As a redox-active metal, copper serves as a catalytic cofactor for critical enzymes such as cytochrome c oxidase, Cu/Zn-superoxide dismutase, and laccases ^6–8^. However, this same redox activity makes excess copper highly toxic, primarily through the generation of reactive oxygen species (ROS), which inflict damage on lipids, proteins, and nucleic acids ^9^. Over the past decade, anthropogenic activities including mining, industrial waste discharge, and excessive use of copper-based agrochemicals have led to elevated levels of copper in soil and water, prompting intensive research into its toxicological impacts on plant growth and cellular function^10^. Numerous studies have documented the adverse effects of copper toxicity on seed germination, root and shoot elongation, chlorophyll biosynthesis, and photosynthetic efficiency ^11–13^. At the cellular level, copper induces oxidative bursts, resulting in ROS accumulation, membrane disruption, protein oxidation, and DNA damage ^14^. These stressors trigger a suite of antioxidant responses involving superoxide dismutase (SOD), catalase, and ascorbate peroxidase ^15^.

Despite extensive research on the physiological, biochemical, and even mitochondrial responses to copper toxicity, very limited attention has been given to the nucleus the master regulator of stress-responsive gene expression and to chromatin-level changes in plants under copper stress ^16^. While drought, salt, and other abiotic stresses are known to influence nuclear morphology, chromatin compaction, and histone modifications, similar investigations under heavy metal stress, especially copper, remain scarce ^17^. In animal, copper has been implicated in mitochondrial dysfunction, cytochrome c release, and redox imbalance, but nuclear-level disruptions remain underexplored ^18^. A few studies on metals like cadmium and aluminum have shown that heavy metals can influence nucleosome positioning, histone methylation, and global transcription patterns, yet comparable mechanisms under copper stress particularly in agriculturally relevant species like *Solanum lycopersicum* have not been systematically investigated ^19,20^. Although copper’s uptake, translocation, and detoxification in plants have been well studied, the downstream nuclear consequences, especially alterations in chromatin organization and nuclear architecture, remain poorly understood^21^. This creates a major gap in our understanding of how copper-induced stress is perceived and transduced at the genomic level, where transcriptional reprogramming is ultimately orchestrated.

This study aims to fill this critical knowledge gap by focusing on nuclear structure and chromatin remodeling as central mediators rather than passive endpoints of copper-induced stress responses^22^. We utilized the root apex of tomato (*Solanum lycopersicum*) as a model due to its sensitivity to environmental cues and its well-defined zones of cellular activity. By combining high-resolution imaging with organelle-specific fluorescent markers and immunostaining techniques, we evaluated how increasing doses of copper affect nuclear morphology and chromatin distribution^23^. To visualize these subcellular changes with precision, we employed confocal microscopy for volumetric organelle imaging and Super-Resolution Radial Fluctuations (SRRF) to resolve fine chromatin structures and nuclear envelope details beyond the diffraction limit^24^. To establish a mechanistic link between copper exposure and nuclear responses, we simultaneously examined mitochondria (for structural damage, membrane potential, cytochrome c release), ROS accumulation, lysosomal activity (autophagy), and energy/redox signaling markers (AMPK, mTOR, NRF2)^25–27^. These multi-organelle indicators allowed us to corroborate that chromatin rearrangement is a direct consequence of systemic copper-induced stress and not merely a secondary or isolated nuclear phenomenon.

We observed profound nuclear alterations in root tip cells under copper stress, including nuclear shrinkage, lobulation, chromatin condensation, and peripheral heterochromatinization all indicative of suppressed transcriptional activity. These nuclear changes occurred in parallel with mitochondrial dysfunction, ROS buildup, and activation of lysosome-dependent autophagy pathways. Additionally, activation of AMPK, inhibition of mTOR, and nuclear translocation of NRF2 pointed to disrupted energy homeostasis and redox signaling. Taken together, these findings support the central role of the nucleus as an integrator of environmental stress signals, rather than a passive responder. Our results not only highlight nuclear architecture as an early and sensitive indicator of copper toxicity but also open new avenues for developing chromatin-based biomarkers for environmental monitoring and stress-resilient crop breeding.

## Results

We cultivated *Solanum lycopersicum* under controlled conditions and exposed them to varying concentrations of copper for 24 hours to induce differential concentration. To assess subcellular effects, we used confocal and SRRF imaging with organelle- and nucleus-specific fluorescent probes and immunostaining. MitoTracker Green (MTG) and TMRE were used to evaluate mitochondrial morphology and membrane potential, while cytochrome c immunostaining detected its release into the cytosol, indicating apoptosis initiation^28–30^. Lysosomal activity was tracked with LysoTracker Red (LTR)^31^. Immunostaining for AMPK and mTOR revealed energy stress signaling and non-canonical nuclear mTOR accumulation^32^. NRF2 immunostaining demonstrated ROS-induced nuclear translocation^33,34^. Nuclear integrity and chromatin remodeling were assessed via DAPI, Hoechst, and H3K4me3 staining^35–37^. PI staining marked late-stage cell death^38^. This integrative approach enabled high-resolution mapping of copper-induced organelle dysfunction, nuclear changes, and cell fate in tomato root apex cells. Additionally, we have checked physiological changes in the root and shoot system **(Supplementary Figures 1)**^39^.

### 1. Early Mitochondrial Collapse and Pro-Apoptotic Signaling Triggered by Copper Exposure

We analyzed mitochondria because they are the primary sites of copper-induced oxidative stress and energy disruption^40^. Early issues with mitochondria are a big reason for stress signals in the cell, which include making reactive oxygen species (ROS), starting cell death (apoptosis), and changes in the cell’s nucleus^41–43^. Exposure to increasing concentrations of copper led to progressive deterioration of mitochondrial integrity in tomato root apex cells, marking the first detectable signal of cellular distress.

MitoTracker Green (MTG) staining showed a clear loss of the typical mitochondrial network, with dysfunctional and fading signals as copper levels increased **(Figure 1 Ia-f)**^28^. This was accompanied by a noticeable drop in mitochondrial membrane potential, shown by the decrease in TMRE fluorescence^30^. In control cells, the signals from MTG and TMRE were evenly spread throughout the root meristem, showing that the mitochondria were healthy and functioning properly **(Figure 1 Ia)**. However, as copper levels increased, the TMRE signal decreased much earlier than the MTG signal, indicating that the mitochondria start to malfunction before they lose their structure **(Figure 1 Ib-f)**. At the same time, cytochrome c (CytC) was found in small spots linked to mitochondria in control samples, but it spread out and moved into the cytoplasm, especially with moderate to high copper exposure **(Figure 2 Ia-f)**^29^. This release of CytC into the cytoplasm fluid is a key sign that the outer membrane of the mitochondria is breaking down and that the process of cell death is starting. Line profile analyses further validated the spatial separation of MTG and CytC signals under stress conditions, confirming that CytC dissociates from fragmented mitochondrial compartments **(Figure 1 & 1&2 IV-V)**. Quantitative measurements revealed a consistent decrease in MTG and TMRE signal intensity, coupled with a significant rise in CytC fluorescence, establishing a strong correlation between mitochondrial breakdown and early apoptotic signaling **(Figures 1 & 2 II-III)**. These findings demonstrate that mitochondrial dysfunction, manifested as membrane depolarization and cytochrome c release, is the first physiological disturbance triggered by copper toxicity, setting off the downstream cascade of stress responses^27^. This event is the first signal that something is wrong, acting as the primary intracellular alarm leading for copper stress to nuclear and cellular remodeling.

**Figure 1.**
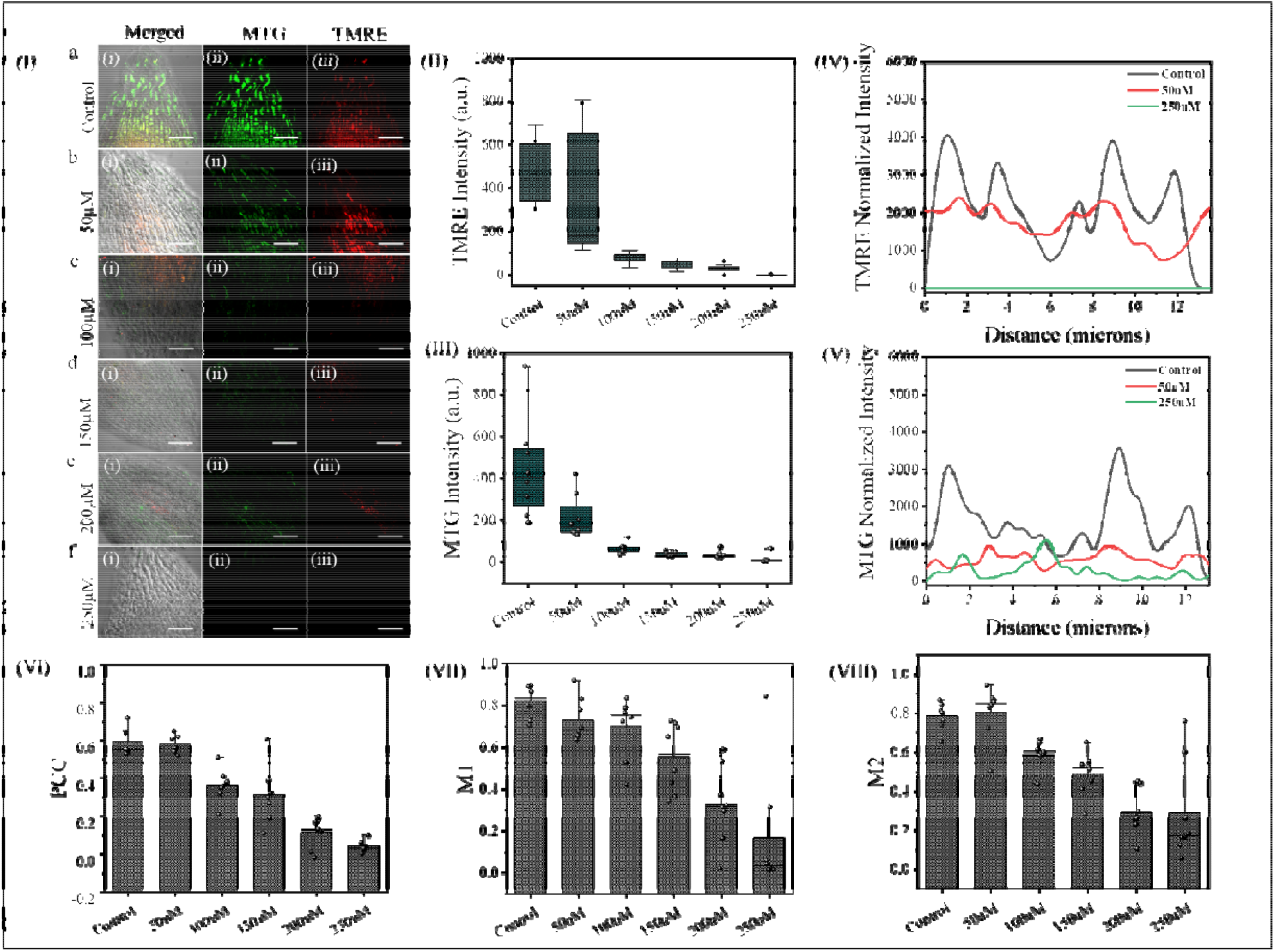
Copper stress leads to mitochondrial membrane depolarization in tomato root apex cells. **(I)** Confocal images show MTG (green) and TMRE (red) staining across increasing copper concentrations. Control cells display strong TMRE signal co-localized with MTG, indicating healthy, polarized mitochondria. Under copper stress, TMRE signal diminishes progressively, while MTG signal remains, indicating loss of membrane potential. **(II–III)** Quantification shows a dose-dependent decrease in TMRE intensity with relatively stable MTG levels. **(IV–V)** Line profiles confirm reduced TMRE signal relative to MTG under stress. (VI–VIII) Bar graphs show reduced mitochondrial area, TMRE-positive cells, and TMRE/MTG ratio, confirming early mitochondrial depolarization under copper stress. Scale bars: 50□µm. Data represent mean ± SEM, n = 10 root tips per condition.

**Figure 2.**
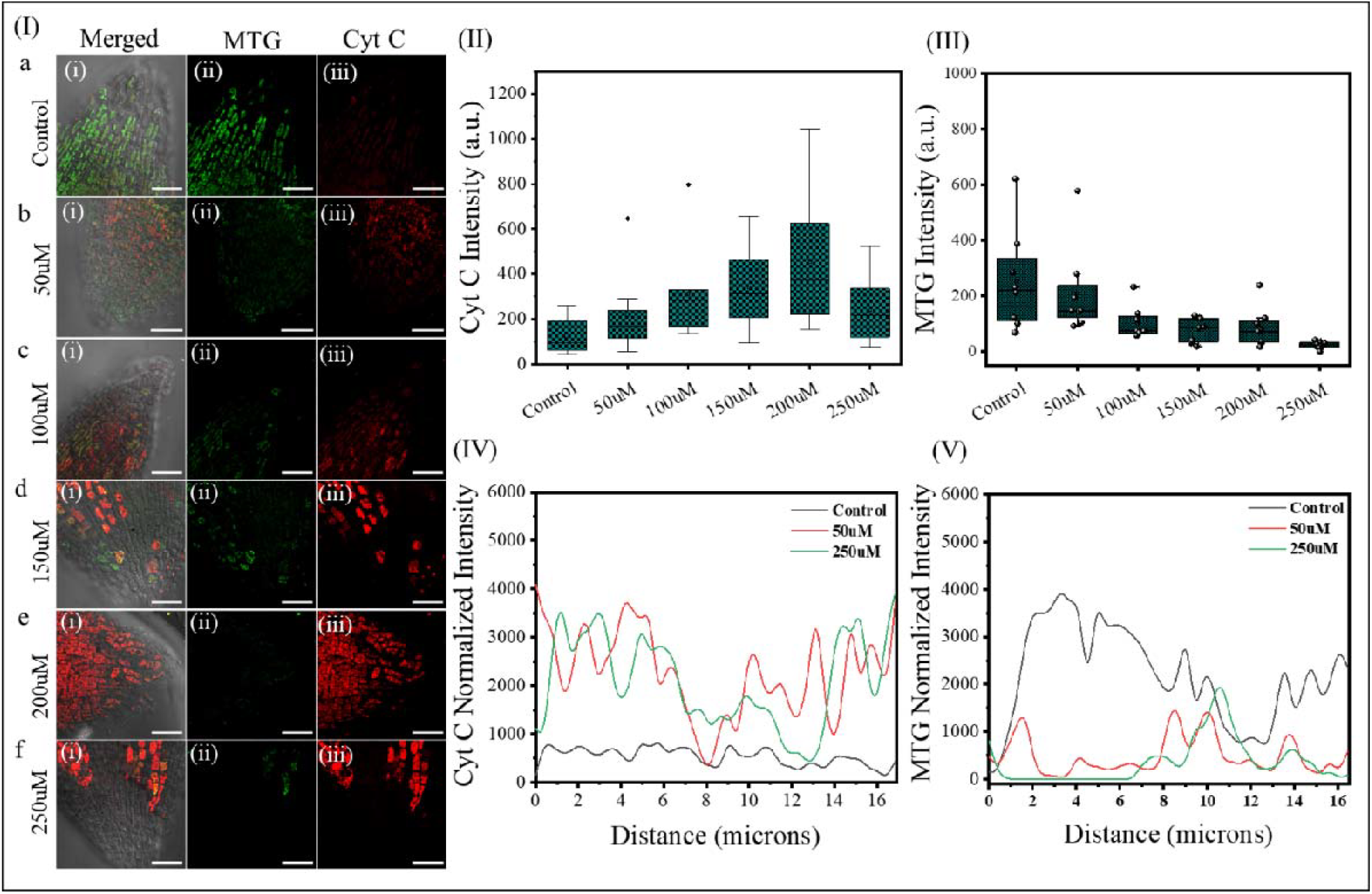
Copper stress triggers mitochondrial damage and cytochrome c release in tomato root apex cells. **(I)** Confocal images show dose-dependent mitochondrial dysfunction (MTG, green) and increased cytochrome c release (Cyt C, red) from mitochondria into the cytosol. **(II–III)** Quantification reveals a rise in Cyt C intensity and a decline in MTG signal with increasing copper concentration. **(IV–V)** Line profiles confirm Cyt C displacement and MTG signal loss, indicating early mitochondrial dysfunction and initiation of apoptotic signaling. Scale bars: 50□µm. Data represent mean ± SEM, n = 10 root tips per condition.

### 2. Mitochondrial Dysfunction Leads to ROS Accumulation and NRF2-Mediated Nuclear Signaling

After mitochondrial dysfunction, exposure to copper caused a significant rise in reactive oxygen species (ROS) levels in the tip cells of tomato roots, showing that mitochondria are a key source of oxidative burst^42^. As shown in **Figure 3**, ROS-specific fluorescent dye staining revealed minimal fluorescence in control cells^34^. The signal became stronger as the amount of copper increased, and measurements showed a significant rise in ROS levels with higher copper concentrations. Line profile analysis showed how ROS is spread out, with high intensity peaks in areas experiencing strong mitochondrial stress, suggesting that damaged mitochondria are the main source of ROS production **(Figure 3 III)**. This ROS surge likely contributes to oxidative damage to lipids and proteins, but more critically, it acts as a secondary messenger, activating downstream redox-sensitive pathways^9^.

**Figure 3.**
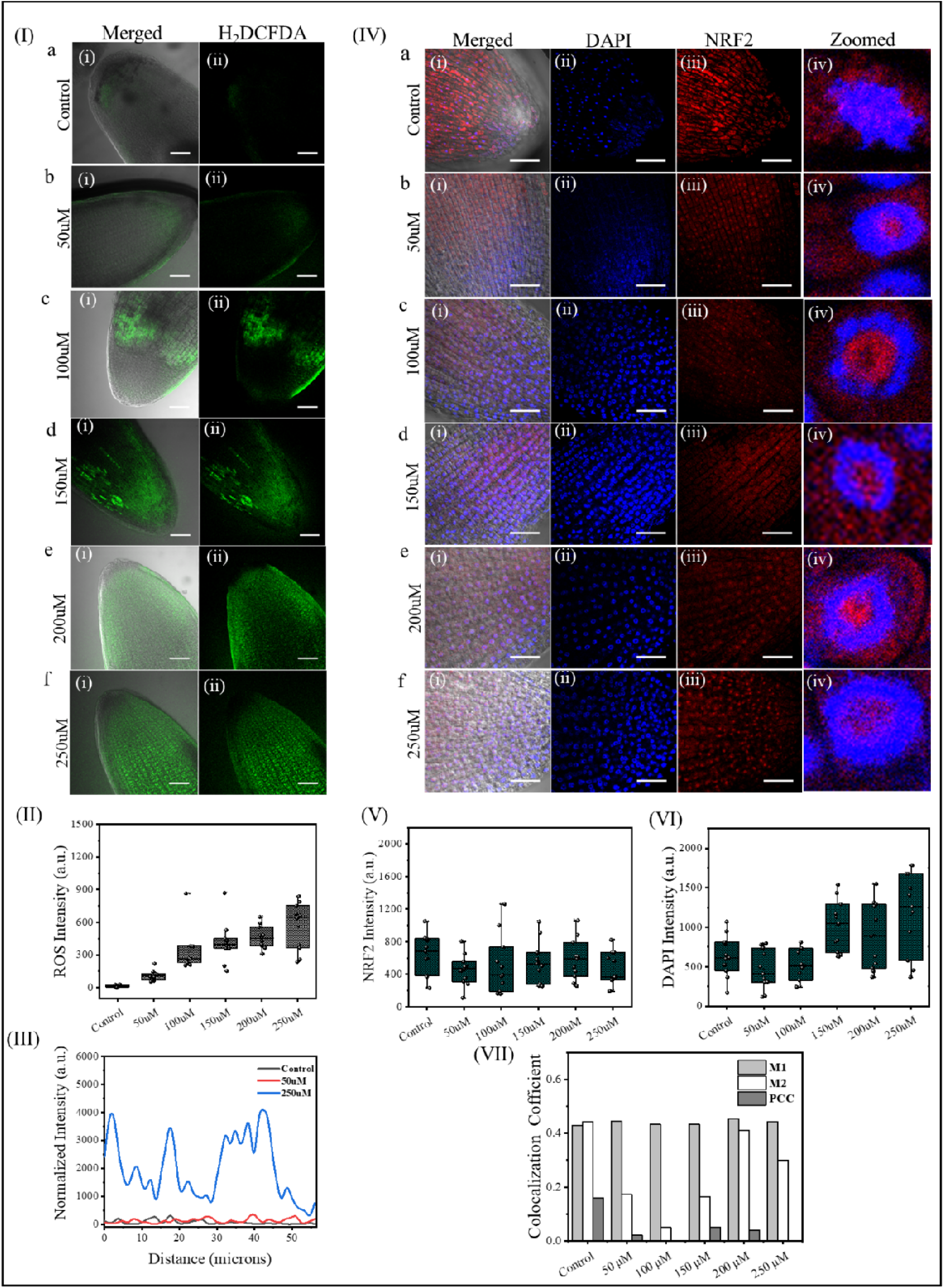
Copper stress induces ROS accumulation and NRF2 nuclear translocation in tomato root apex cells. **(I)** H□DCFDA staining shows a dose-dependent increase in ROS levels under copper treatment. **(II–III)** Quantification and line profile confirm elevated ROS, especially at 250□µM. **(IV)** Confocal images show NRF2 translocation from cytoplasm to nucleus, confirmed by DAPI co-staining and zoomed views. **(V–VII)** Box plots and co-localization analysis (M1, M2, PCC) support increased nuclear NRF2 accumulation and chromatin condensation. Scale bars: 50□µm & 2□µm. Data: mean ± SEM, n = 10 root tips.

The transcription factor NRF2 mediates one such response, and **Figure 3 IV** uses immunofluorescence to visualize its activation under oxidative stress^33^. In control cells, NRF2 exhibited a weak, diffuse cytoplasmic signal with minimal nuclear localization **(Figure 3 IVa)**. NRF2 fluorescence became noticeably nuclear-enriched after being treated with copper, especially in the early root apex zones. High-resolution insets and DAPI co-staining confirmed NRF2 translocation into the nucleus **(Figure 3 IVb-f)**^36^. This change shows the typical redox regulation process, where ROS help keep NRF2 from breaking down, letting it build up and move into the nucleus^44^. Quantitative analysis revealed a significant increase in the nuclear-to-cytoplasmic ratio of NRF2 signal across copper concentrations **(Figure 3 V-VI)**. These results indicate that oxidative stress connects damage to mitochondria with the cell’s response in the nucleus, with NRF2 playing an important role in helping the cell adapt by turning on stress-related genes. This translocation likely initiates protective antioxidant gene expression while also priming the nucleus for broader chromatin remodeling in response to sustained cellular damage.

### 3. Copper-induced oxidative stress reprograms redox-energy signaling by activating AMPK and promoting the non-canonical nuclear accumulation of mTOR

To investigate the metabolic response to oxidative stress triggered by copper, we examined the activation of AMP-activated protein kinase (AMPK) and its downstream target mTOR in tomato root apex cells ^45,46^. As shown in **Figure 4**, immunostaining of phosphorylated AMPK (pAMPK) revealed a low basal level in control cells, where fluorescence remained diffuse and weak across the cytoplasm **(Figure 4 Ia)**^32^. Upon exposure to increasing copper concentrations, the pAMPK signal intensified significantly and became enriched in both the cytoplasm and nucleus, indicating activation under energy-deprived conditions **(Figure 4 Ib-f)**. Quantitative analysis confirmed a strong dose-dependent increase in pAMPK intensity, reflecting cellular energy stress and metabolic reprogramming **(Figure 4II-V)**.

**Figure 4.**
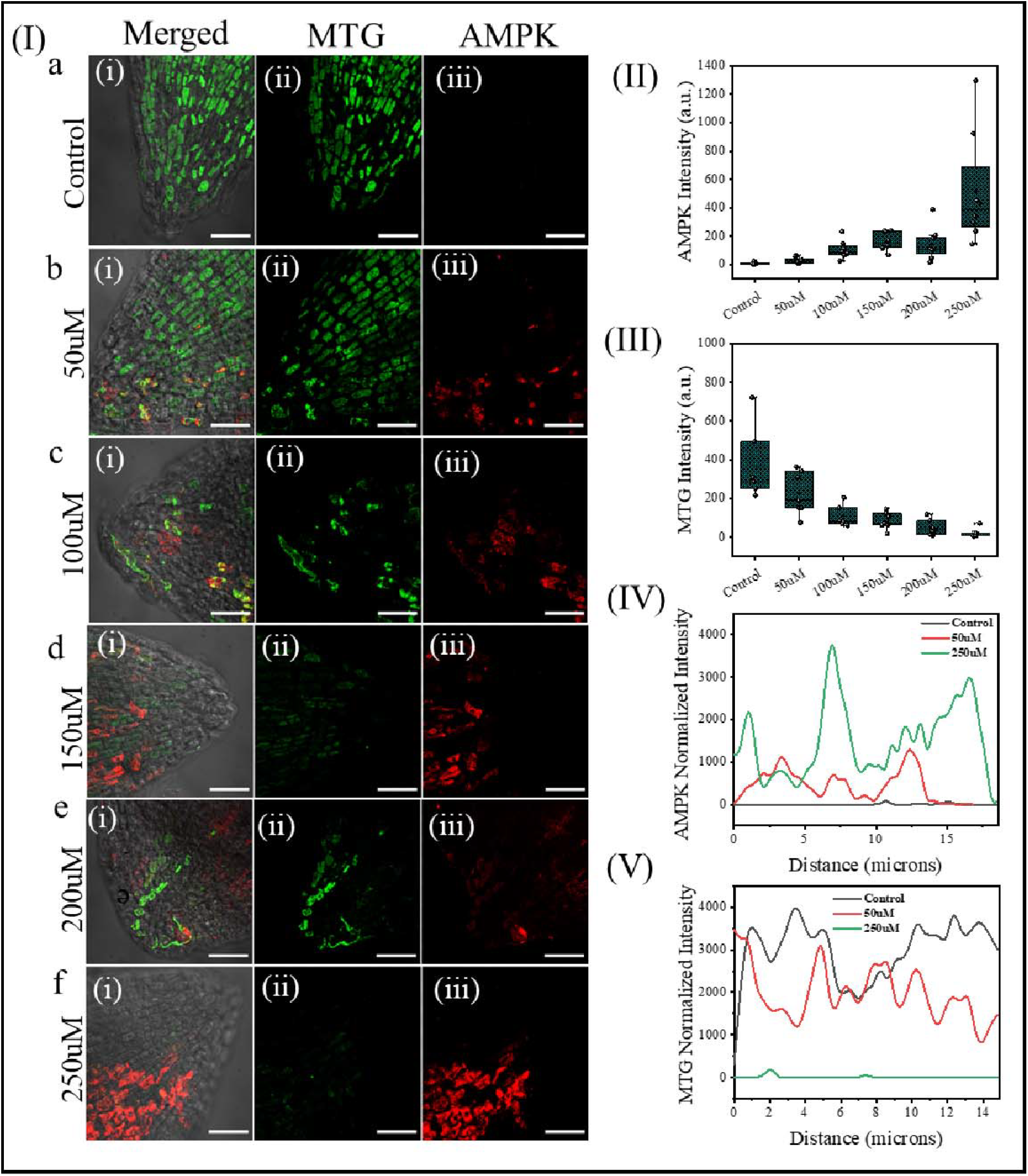
Copper stress activates AMPK and disrupts mitochondrial integrity in tomato root apex cells. **(I)** Confocal images show mitochondrial structure (MTG, green) and AMPK (red) localization under increasing copper concentrations. **(II–III)** Quantification reveals dose-dependent AMPK activation and a decline in MTG signal. **(IV–V)** Line profiles confirm increased AMPK intensity and reduced mitochondrial mass under stress, indicating energy sensing activation coupled with mitochondrial dysfunction. Scale bars: 50□µm. Data represent mean ± SEM, n = 10 root tips per condition.

In parallel, we observed a marked suppression of mTOR expression under copper treatment **(Figure 5)**. In control root cells, mTOR signal was predominantly cytoplasmic and enriched in actively dividing regions **(Figure 5Ia)**. However, with increasing copper exposure, cytoplasmic mTOR intensity progressively declined **(Figure 5Ib-f)**. Interestingly, and in contrast to canonical mTOR localization, we observed a consistent and copper dose-dependent nuclear accumulation of mTOR, as visualized in high-resolution optical sections and confirmed in supplementary data **(Supplementary Figure S2)**. This movement of mTOR into the nucleus might indicate a different role for mTOR in managing stress-related gene activity or changes in DNA structure, which has been suggested in animals but not yet studied in plants^47^. Line profile analysis across nuclear boundaries further validated the spatial redistribution of mTOR from cytoplasm to nucleus under stress **(Figure 5II-V)**. Taken together, these findings suggest that copper-induced oxidative stress activates AMPK and suppresses mTOR-mediated anabolic signaling^46^. Additionally, the surprising presence of mTOR in the nucleus during stress suggests it might play a role in controlling gene activity or changes in gene expression, possibly connecting how the cell responds to stress with its energy use. These results show that the AMPK–mTOR pathway is a key connection between oxidative stress, energy problems, and changes in the cell’s nucleus during heavy metal toxicity.

**Figure 5.**
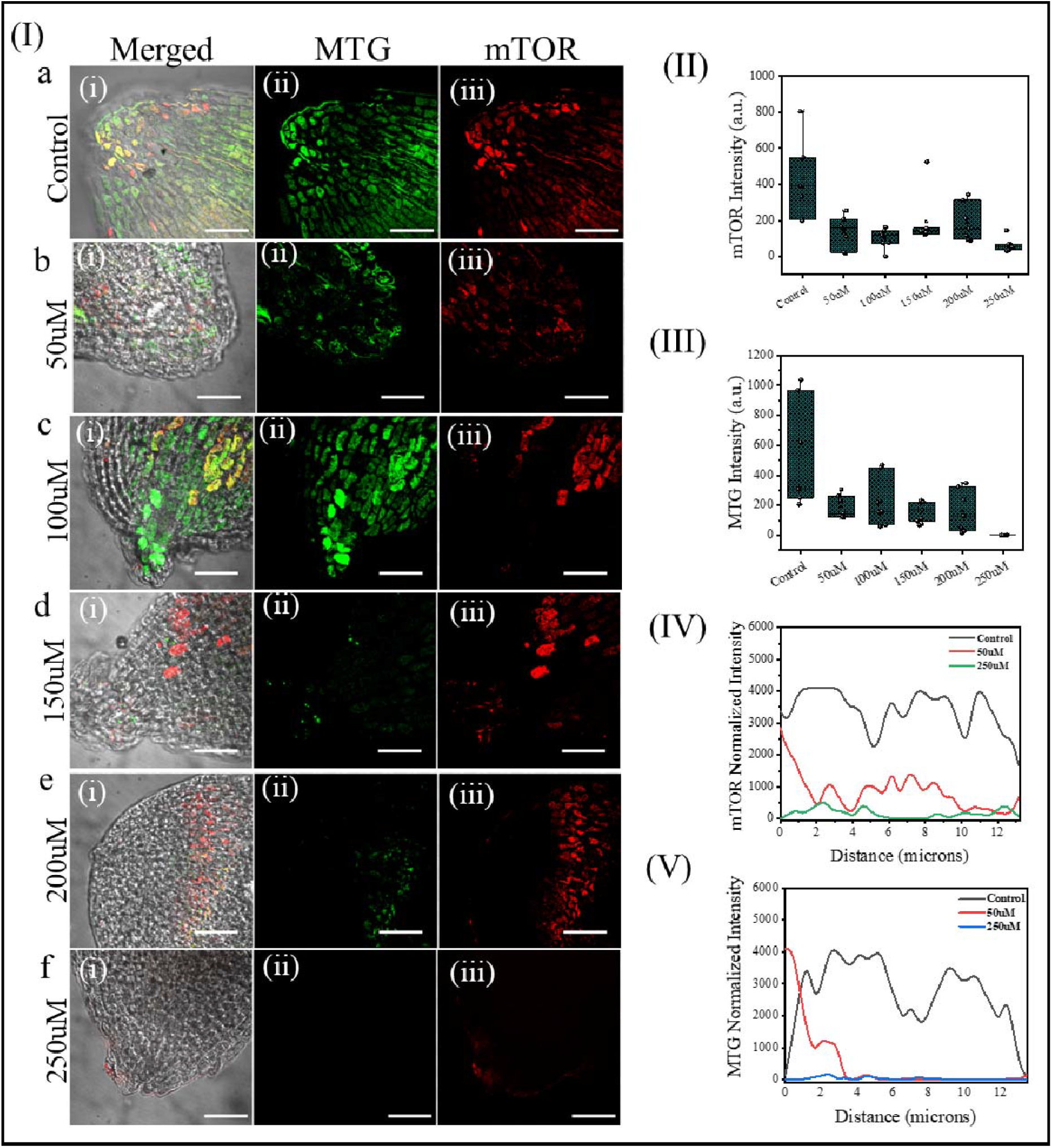
Copper stress reduces mTOR expression and mitochondrial integrity in tomato root apex cells. **(I)** Confocal images show MTG (green) and mTOR (red) under increasing copper concentrations. Control cells show strong cytoplasmic mTOR and intact mitochondria; copper stress causes reduced mTOR signal and mitochondrial fragmentation. **(II–III)** Box plots show a dose-dependent decline in mTOR and MTG intensity. **(IV–V)** Line profiles confirm spatial reduction in mTOR and mitochondrial signal with increasing copper levels. Scale bars: 50□µm. Data represent mean ± SEM, n = 10 root tips.

After activating AMPK and reducing mTOR due to copper stress, we looked at autophagosomes formation as we already see that the mitochondria getting damaged or dysfunctioned. Lysotracker Red (LTR) staining revealed a progressive increase in lysosomal activity in tomato root apex cells with increasing copper concentrations **(Supplementary Figure 3)**^31^. In control cells, lysosomes appeared small and sparsely distributed, while copper treatment led to enlarged, intensely labeled vesicles localized near fragmented mitochondria, suggesting mitophagy induction (**Supplementary Figure 3 Ia-f)**. Intensity profile analysis confirmed lysosomal intensity increases with increase in copper concentration especially under moderate copper levels **(Supplementary Figure 3 II)**. These changes occurred downstream of AMPK activation, consistent with mTOR inhibition relieving autophagic suppression. However, when there is too much copper, lysosomes group together too much, which might mean they are overwhelmed or stressed, possibly leading to problems with the nucleus and changes in the structure of DNA that are noticed later on. Collectively, these findings indicate that lysosomes initially mitigate damage, but under prolonged stress, they may become dysfunctional, shifting the balance from survival to programmed cell death.

### 4. mTOR–NRF2 Signaling Orchestrates Peripheral Chromatin Re-Arrangement and Progressive Condensation under Copper Stress

Copper-driven oxidative and metabolic stress culminated in pronounced reorganization of nuclear architecture using DAPI and Hoechst **(Supplementary Figure 4)** in tomato root apex cells **(Figure 6)**^36^. High-resolution DAPI images show that nuclei in control roots are large, evenly stained, and euchromatin-rich **(Figure 6Ia)**. With increasing copper, nuclei first adopt a “ring-like” pattern in which chromatin relocates to the nuclear periphery while the central region appears partially cleared **(Figure 4Ib)**^48^. Quantification confirmed a progressive decline in nuclear cross-sectional area and intensity of the nucleus **(Figure 6 II-III)**. Centre of mass analyses reveal that peripheral chromatin density rises first, followed by a global spike in total chromatin signal once full condensation is reached **(Figure 6 IV)**.

**Figure 6.**
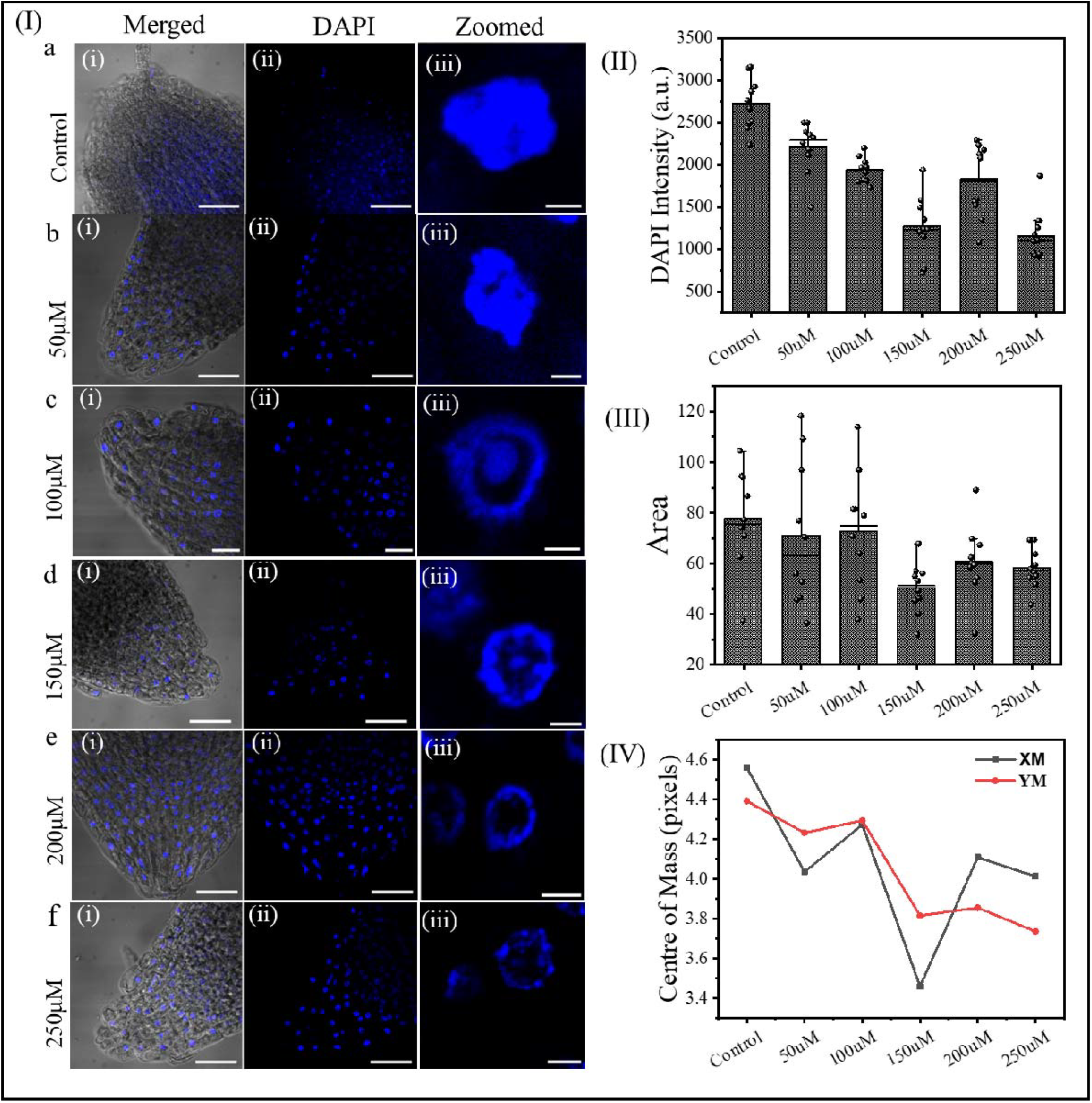
Copper stress induces nuclear arrangement and displacement in tomato root apex cells. **(I)** Confocal images of DAPI-stained nuclei show progressive chromatin condensation and nuclear shrinkage under increasing copper concentrations, with control cells displaying large, round euchromatic nuclei. **(II–III)** Quantification shows reduced DAPI intensity and nuclear area with rising copper, indicating chromatin compaction. **(IV)** Centre of mass analysis reveals displacement of nuclear position, suggesting spatial reorganization under stress. Scale bars: 50□µm and 2□µm. Data are mean ± SEM, n = 10 root tips.

These structural changes coincide spatiotemporally with the nuclear accumulation of both mTOR and NRF2 **(shown in Figures 5–6 and Supplementary S2)**, two factors known to recruit or modulate histone-modifying enzymes. Stress-activated mTOR has been found to work with chromatin remodellers, while nuclear NRF2 helps activate antioxidant genes by changing how easily nucleosomes can be accessed^49^. Their simultaneous presence in the nucleus under copper stress plausibly drives the observed transition, initial peripheral tethering of chromatin (a reversible, gene-regulatory stage) followed by irreversible global compaction when cellular recovery fails. Thus, the nucleus is not merely a passive casualty, it actively remodels its chromatin landscape in response to converging metabolic (mTOR) and redox (NRF2) cues, ultimately locking the cell into a condensed, transcriptionally silent state that precedes programmed cell death.

### 5. Chromatin Condensation Leads to Terminal Commitment to Cell Death

To study how chromatin changes during stress from copper, we used a technique called immunostaining to look for H3K4me3, a marker that shows active gene areas in the chromatin^35^. In control tomato root apex cells, the H3K4me3 signal was evenly distributed throughout the nucleus and chromatin are thread like, indicating a transcriptionally permissive chromatin state and no condensation **(Figure 7 Ia)**^50,51^. With increasing copper concentrations, the H3K4me3 signal shifted toward the nuclear periphery, forming ring-like patterns and chromatin is condensed to the periphery, as we see the intensity of the chromatin was higher at periphery **(Figure 7 Ib--e)**. At higher copper doses, this peripheral localization gave way to strong nuclear compaction, reflecting the collapse of euchromatic architecture and it comes again in the centre **(Figure 7 If)**. We quantitatively assessed these chromatin changes by measuring the domain number and cross-sectional area using fluorescence intensity plots and eccentricity **(Figure 7 II-IV)**. The number of domains and the area they cover showed that the chromatin moved to the edges, and the eccentricity indicated a loss of euchromatin features, which matches with reduced gene activity.

**Figure 7.**
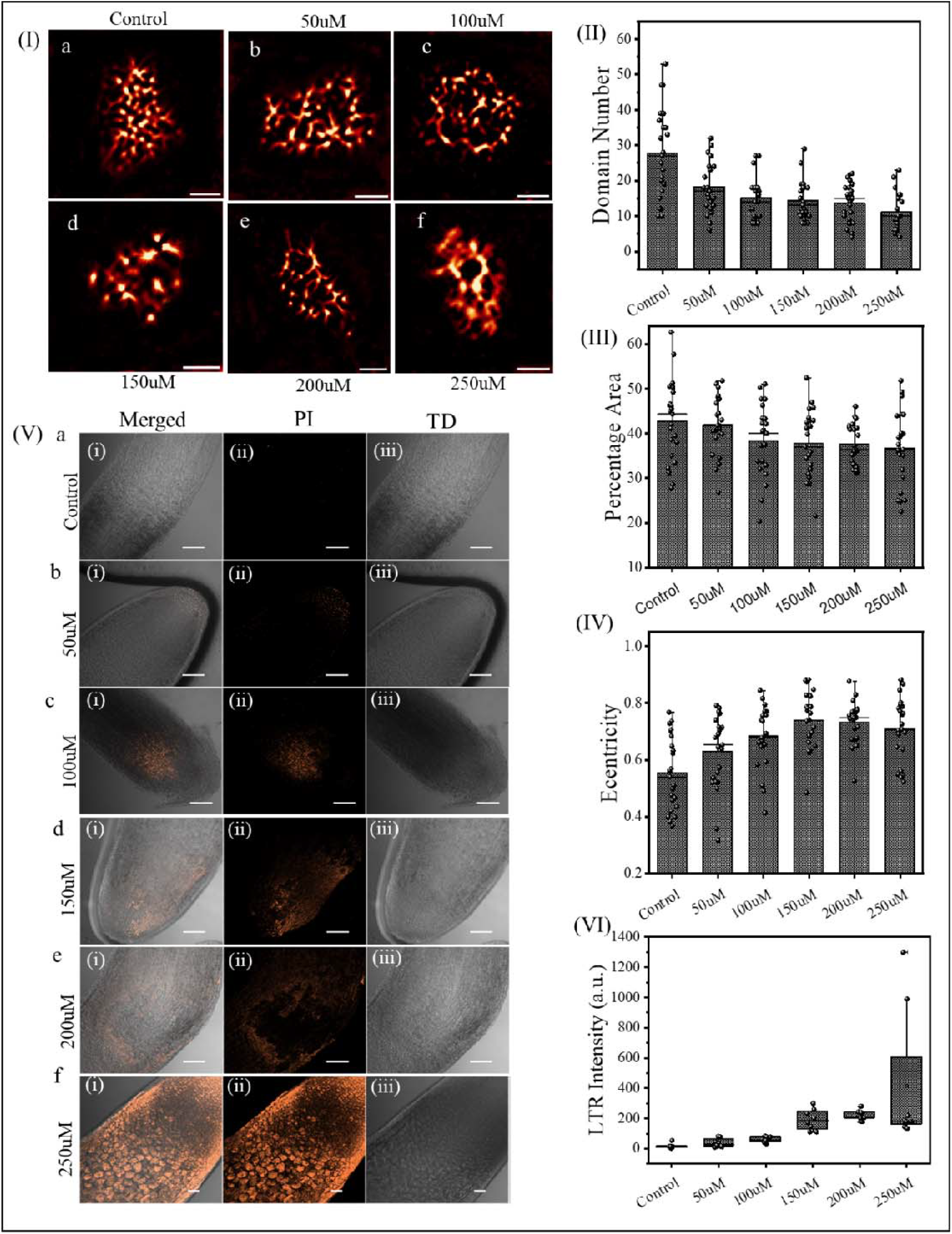
Copper stress promotes euchromatin compaction and cell death activation in tomato root apex cells. (I) SRRF super-resolution images of H3K4me3 staining reveal a decline in euchromatin domain number and area with increasing copper. (II–IV) Quantification shows a decrease in euchromatin domain number, area, and an increase in domain eccentricity, indicating chromatin compaction. (V) Confocal images of PI staining show dose-dependent increases in membrane-compromised cells and acidic vesicles. (VI) PI fluorescence intensity increases significantly at 250□µM, suggesting cell death. Scale bars: 2□µm (I), 50□µm (V). Data represent mean ± SEM, n = 10 root tips.

To see how much cell death happened due to copper stress, we used propidium iodide (PI) staining on tomato root tip cells^38^. In the control samples, the PI signal was almost not seen, which means the plasma membranes were intact and the cells were alive **(Figure 7 Va)**. However, with increasing copper concentrations, a progressive and region-specific accumulation of PI signals was observed, particularly in the elongation and differentiation zones of the root apex **(Figure 7 Vb-d)**. At higher copper doses, the PI signal became intense and widespread, indicating extensive loss of membrane integrity, a definitive marker of late-stage cell death **(Figure 7 Ve-f)**^52^. The PI-positive staining shows a step-by-step process of early and late cell death, which matches earlier findings of mitochondrial cytochrome c release and chromatin condensation. Fluorescence demonstrated spatial correlation between PI-positive cells and nuclear collapse, confirming that these death signals originate from stress-compromised regions **(Figure 7 VI)**. These findings demonstrate that copper exposure compromises membrane integrity and triggers irreversible cell death once the cell surpasses its compensatory thresholds. This evidence supports the conclusion that cell death under copper stress is a downstream consequence of mitochondrial collapse, oxidative damage, and nuclear disintegration.

### 6. Shoots Exhibit Delayed Chloroplast and Nuclear Alterations Compared to Roots

As this data demonstrates that copper toxicity affects the shoot and root systems differently in tomato plants. In the shoot, chloroplast autofluorescence remains largely intact at lower Cu^2^□ concentrations (≤100□µM), indicating minimal structural disruption^53^. However, at higher concentrations (≥250□µM), chloroplast morphology becomes irregular and signal intensity decreases, reflecting functional impairment. The co-staining with LysoTracker Red (LTR) shows increasing lysosomal activity and autophagy, confirmed by colocalization in the merged images **(Figure 8I-III)**^31^.

**Figure 8.**
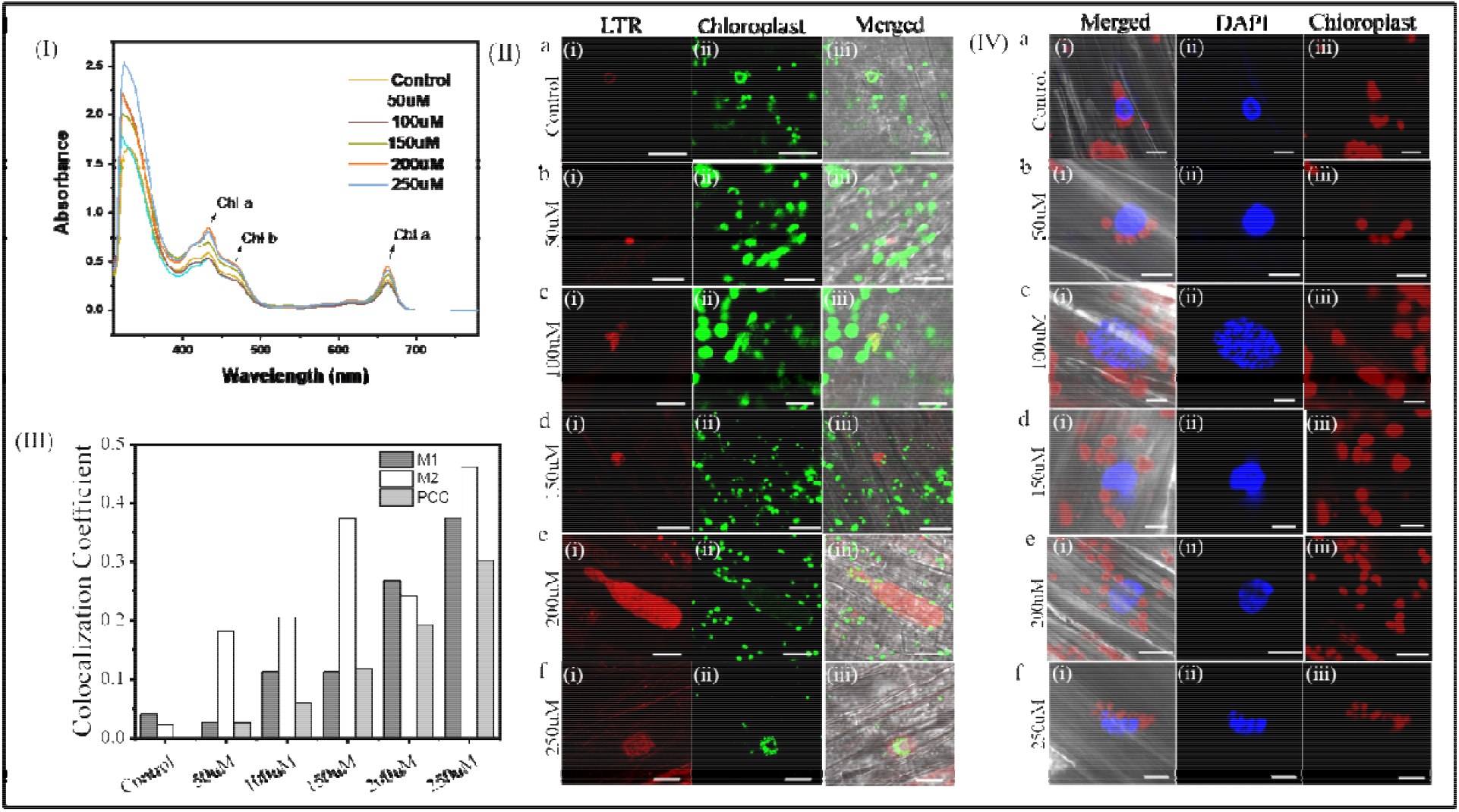
Copper-induced chloroplast and nuclear alterations in tomato root and shoot tissues. (I) UV-vis spectra and bar graph showing pigment reduction with increasing Cu^2^□ concentrations. **(II-III)** Confocal images of shoot cells stained with LTR (red) and chloroplast autofluorescence (green) reveal intact chloroplasts at low Cu^2^□ levels but disruption at higher doses induce autophagy, as seen in colocalization graph. **(IV)** Shoot cells show chloroplast-nucleus association. Scale bars: 50□µm and 20□µm. Data are mean ± SEM, n = 10 root tips.

Despite these changes, the nuclear morphology in shoot cells remains relatively preserved, with nuclei maintaining a central position and showing no significant condensation or peripheral chromatin shift^54^. This contrasts sharply with the root system, where nuclei exhibit shrinkage, lobulation, and chromatin margination even at moderate Cu^2^□ levels^51^. These differences suggest that root cells are more severely and earlier affected by copper stress, likely due to their direct contact with the copper-rich growth medium **(Figure 8 IV)**. Additionally, chloroplast–nucleus association observed in shoot cells may indicate early stress signaling or protective interactions, though without the pronounced nuclear reorganization seen in roots. Overall, the data presented in **Figure 8** support the conclusion that roots are more prone to copper-induced damage, while shoot tissues respond more gradually and maintain compartmental integrity under similar conditions.

## Discussions

Copper toxicity in plants often arises from excessive copper in soil, primarily due to industrial waste, mining activities, and overuse of copper-based fertilizers and pesticides. This leads to the accumulation of free Cu^2^□ ions in plant tissues, causing oxidative stress, organelle damage, enzyme inactivation, and disruption of nutrient balance ultimately impairing plant growth and development. This study provides a detailed look at how copper toxicity affects different parts of the cell in *Solanum lycopersicum*, showing a series of problems with cell structures, stress responses, changes in DNA packaging, and programmed cell death. Our results indicate that mitochondria are the earliest sensors of copper stress, exhibiting fragmentation, membrane depolarization, and cytochrome c release, hallmarks of intrinsic apoptotic signaling^18,27^. This mitochondrial collapse leads to a secondary surge in reactive oxygen species (ROS), which contributes to oxidative damage and serves as a signaling cue for downstream regulatory responses^42^. One important discovery from this study is that NRF2 moves into the nucleus and AMPK gets activated, showing a link between redox and energy stress signals^49^. Unexpectedly, we also observed a non-canonical nuclear localization of mTOR under high copper conditions, suggesting potential involvement in chromatin regulation. These signaling events collectively primed the nucleus for structural reorganization. High-resolution DAPI and H3K4me3 imaging revealed a progressive shift from euchromatin-rich, transcriptionally active nuclei to highly condensed, transcribed silent chromatin states. The initial buildup of chromatin at the edges, followed by overall tightening, shows a step-by-step response of the nucleus to stress that is closely connected to metabolism and redox signaling. However, under sustained copper exposure, excessive lysosomal activity may contribute to nuclear destabilization and eventual cell death^55^. Propidium iodide staining showed that the cell membranes broke down in the later stages, leading to cell death, which matched areas where the genetic material fell apart and the mitochondria stopped working^52^. Taken together, our data reveal that the nucleus acts not merely as a passive target but as an active integrator of organelle-derived stress. The sequential activation of the mitochondria-ROS-AMPK/mTOR-NRF2 chromatin axis under copper toxicity establishes a molecular framework for understanding how environmental stress culminates in programmed cell death. Importantly, the identification of nuclear mTOR localization and chromatin reorganization as early stress markers enables the development of targeted strategies to enhance metal stress tolerance in plants **(Figure 9)**.

**Figure 9.**
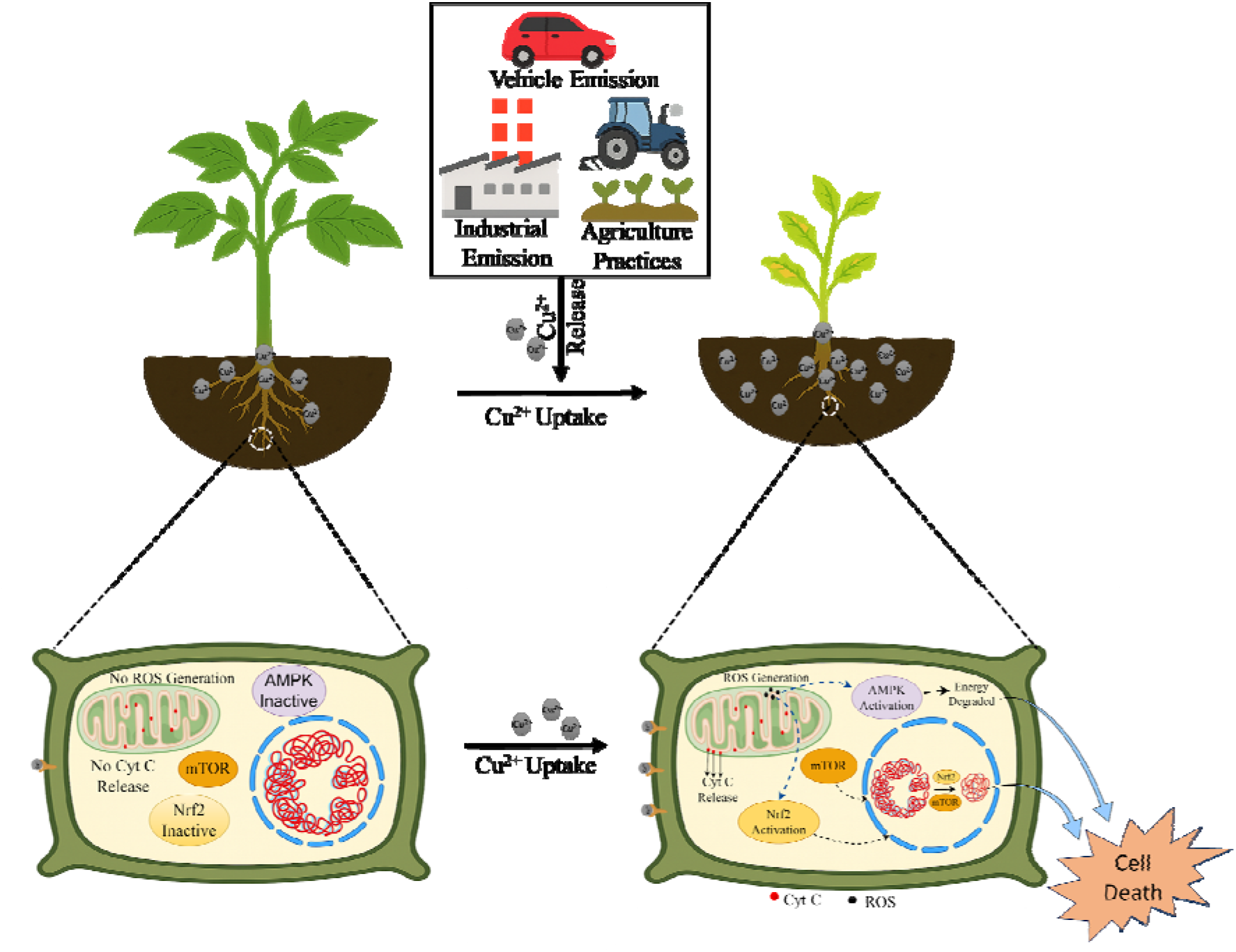
Copper-induced cellular stress in plant roots. Under normal conditions (left), cellular pathways remain inactive with no ROS or Cyt C release. Under copper toxicity (right), excess Cu^2^□ from industrial and agricultural sources triggers ROS production, Cyt C release, AMPK activation, and Nrf2 translocation, ultimately leading to cell death.

We observed that the root tissues were more susceptible to copper-induced stress compared to the shoots. At lower copper concentrations, the shoot cells did not exhibit significant alterations in chloroplast or nuclear dynamics. However, at higher concentrations, both organelles showed noticeable stress responses, indicating that shoots require a higher threshold of copper exposure to exhibit subcellular damage. This suggests a differential sensitivity between root and shoot tissues, with the root acting as the primary site of copper toxicity. This study provides detailed strategies for metal detoxification in plants using multiple complementary approaches, including physiological analysis, organelle-specific imaging, and stress pathway modulation to understand and mitigate copper-induced toxicity.

## Conclusion

This study reveals that copper toxicity in *Solanum lycopersicum* triggers a highly coordinated cascade of organelle dysfunction, redox imbalance, and nuclear remodeling. Mitochondria emerge as the primary sensors of Cu-induced stress, undergoing early membrane depolarization and cytochrome c release. This initiates ROS accumulation and the activation of stress-responsive pathways, notably AMPK and NRF2. We identify a novel nuclear localization of mTOR under copper stress, suggesting its potential role in chromatin dynamics. High-resolution chromatin imaging showed a progressive peripheral relocation and condensation of chromatin, coinciding with transcriptional repression and cell death. These nuclear changes are not passive but actively integrate organelle-derived signals, particularly those related to energy and redox status. Lysosomal activation initially supports cellular survival but becomes overwhelmed under high Cu stress, contributing to nuclear destabilization and apoptosis. Together, these findings provide a mechanistic model for copper-induced programmed cell death, mediated through a mitochondrion–ROS–AMPK/mTOR–NRF2–chromatin axis. Importantly, nuclear architecture and chromatin organization emerge as early and sensitive markers of heavy metal stress, offering potential targets for genetic or chemical interventions aimed at improving plant resilience to environmental contaminants.

## Data and Materials Availability

Most of the data we provided in the main manuscript are available with the manuscript itself and its supplementary information.

## Acknowledgements

The authors thank the Advanced Materials Research Centre (AMRC) and Indian Institute of Technology Mandi (IIT Mandi) for providing the facilities and the sophisticated instruments. All the contributing authors thank the Ministry of Education (MoE), India, for the research scholarship.

## Author Contributions

SC and SHC jointly designed and conceptualized the experiments with input from CKN. SC optimized the protocols and performed all plant experiments, imaging and result analysis. SHC contributed to all experimental procedures, and result analysis. AS assisted with imaging. SC and SHC wrote the manuscript with guidance from CKN. CKN supervised and provided overall direction for the project.

## Ethical and Consent

Ethical approval was not required for this study, as it involved standard physiological experiments on *Solanum lycopersicum* (tomato) root apex cells under copper-induced stress conditions. No genetically modified organisms (GMOs), gene editing, or transgenic lines were used. The experimental procedures complied with institutional and national guidelines for research involving plants.

## Notes

### Competing Interest Statement

The authors have declared no competing interest.

