## Supplementary Data for "Copper Stress Trigger Organelles Communication and Chromatin Condensation Leading to Cell Death in *Solanum lycopersicum*"

**Materials and Methods**

**Plant Growth and Copper Treatment**

*Solanum lycopersicum* (tomato) seeds were surface-sterilized and germinated on half-strength Murashige and Skoog (½ MS) agar plates under controlled growth conditions (25 ± 2 °C, 16/8 h light/dark photoperiod). After two weeks, uniform seedlings were transferred to fresh ½ MS media supplemented with copper sulfate (CuSO₄·5H₂O) at concentrations of 0 (control), 50, 100, 150, 200, and 250 µM. These concentrations were selected to cover a gradient from sub-toxic to overtly toxic levels of copper. Treatments were applied for 24 hours a duration sufficient to induce early cellular responses without causing irreversible tissue damage prior to imaging and downstream analyses.

**Mitochondrial Staining**

To assess mitochondrial status in *Solanum lycopersicum* root apex cells under copper and anaesthetic stress, two fluorescent dyes were employed: MitoTracker Green (MTG) and Tetramethylrhodamine Ethyl Ester (TMRE). MTG is a cell-permeant dye that binds to mitochondrial proteins irrespective of membrane potential, allowing visualization of total mitochondrial mass. Root tips were incubated with 2 µL/mL MTG in phosphate-buffered saline (PBS) for 40 minutes at room temperature in the dark, followed by gentle PBS washes prior to imaging. In contrast, TMRE is a potentiometric dye that selectively accumulates in active mitochondria in response to their membrane potential (ΔΨm). Loss of TMRE fluorescence indicates mitochondrial depolarization, a key marker of mitochondrial dysfunction. For this assay, root tips were stained with 500 nL/mL TMRE for 20 minutes, washed with PBS, and immediately imaged. The combined use of MTG and TMRE enabled differentiation between changes in mitochondrial content and mitochondrial health, providing a comprehensive evaluation of mitochondrial responses to stress

**Immunostaining and nucleus**

Immunolocalization was performed to assess the subcellular localization and expression dynamics of key stress- and epigenetic-response proteins: Cytochrome c (Cyt C), AMP-activated protein kinase (AMPK), mammalian target of rapamycin (mTOR), nuclear factor erythroid 2–related factor 2 (NRF2), and histone H3 lysine 4 trimethylation (H3K4me3). Root tips from *Solanum lycopersicum* seedlings were first fixed in freshly prepared 4% paraformaldehyde (PFA) in phosphate-buffered saline (PBS, pH 7.4) for 30 minutes at room temperature to preserve cellular structure and epitope integrity. Following fixation, samples were rinsed three times in PBS and then permeabilized using 0.2% Triton X-100 in PBS for 15 minutes to facilitate antibody penetration through the plasma and nuclear membranes.

To reduce non-specific antibody binding, the samples were incubated in blocking buffer containing 3% bovine serum albumin (BSA) in PBS for 1 hour at room temperature. Root tips were then incubated overnight at 4°C with primary antibodies targeting either Cyt C, AMPK, mTOR, NRF2, or H3K4me3 (Abclonal or equivalent source). All primary antibodies were diluted at 1:100 in blocking buffer. For multiplexed staining (e.g., co-localization of nuclear and cytoplasmic markers), care was taken to use compatible host species and fluorophores.

Following overnight incubation, samples were washed three times with PBS containing 0.1% Tween-20 (PBST) to remove unbound primary antibodies. Then, the root tips were incubated with species-specific secondary antibodies conjugated to fluorescent dyes either Alexa Fluor 488 (green) or Alexa Fluor 594 (red)—diluted at 1:600 in blocking buffer, for 1 hour at room temperature in the dark. After secondary incubation, samples were washed again with PBST to remove excess antibodies.

To visualize nuclei and confirm nuclear localization of specific proteins, samples were counterstained with either DAPI (1 µg/mL) or Hoechst 33342 (1 µg/mL) for 10 minutes, followed by final PBS washes. Root tips were then mounted on glass slides using a glycerol-based antifade mounting medium and sealed with coverslips prior to imaging. Fluorescent signals were visualized using confocal laser scanning microscopy, and localization patterns were analyzed to assess stress-induced translocation (e.g., Cyt C from mitochondria to cytosol, NRF2 to nucleus) and chromatin remodeling (e.g., changes in H3K4me3 nuclear domains).

**Lysosome and ROS Detection**

Lysosomal activity in *Solanum lycopersicum* root cells was assessed using two complementary fluorescent probes. 1. LysoTracker Red (LTR): A commercially available, cell-permeable fluorescent dye that accumulates in acidic organelles (such as lysosomes) due to its weakly basic nature. 2. Lab-made Zn-complex probe: A custom-synthesized fluorescent probe with pH-sensitive fluorescence designed to detect acidic compartments, allowing parallel validation of lysosomal dynamics. Roots were incubated with 2 µL/mL LTR in PBS for 30 minutes at room temperature in the dark. Following incubation, excess dye was removed by gentle PBS washes. Imaging was performed using a confocal microscope with excitation at 561 nm and emission captured in the 580–630 nm range.

**ROS Detection**: Root tips were incubated in **10 µM H₂DCFDA** prepared in PBS for 30 minutes at room temperature in the dark. Post-staining, roots were washed gently with PBS to remove excess dye and immediately imaged under 488 nm excitation, with emission collected at 510–540 nm. The increase in green fluorescence intensity served as a direct indicator of ROS accumulation under copper stress, enabling quantitative comparison across treatments.

**Cell Death Assessment**

**Propidium Iodide (PI)** staining was used to identify membrane-compromised cells. PI staining was used to assess cell viability in *Solanum lycopersicum* root apex cells following anaesthesia and copper stress treatments. Root tips were incubated with 500 nL/mL of PI solution for 15 minutes at room temperature in the dark. After incubation, samples were washed thoroughly with phosphate-buffered saline (PBS) to remove unbound dye. Upon entering compromised cells, PI intercalates with DNA and emits bright red fluorescence when excited at 561 nm (TRITC channel). Imaging was performed using confocal microscopy, and PI-positive nuclei were quantified to determine the extent of cell death under different experimental conditions.

**Imaging and Microscopy**

1. **Confocal laser scanning microscopy** was performed using a Nikon Eclipse Ti inverted microscope, and image acquisition was conducted with Nikon NIS-Elements software. Excitation/emission wavelengths were adjusted per dye: MTG (488/510–540 nm), TMRE (561/580–620 nm), DAPI/Hoechst (405/440–480 nm), PI (561/600–650 nm), and Alexa-conjugated immunostains (488 or 594).
2. **Super-Resolution Radial Fluctuations (SRRF)** is a computational super-resolution technique that enables the reconstruction of sub-diffraction limit structures from conventional fluorescence microscopy data. SRRF operates by analyzing temporal intensity fluctuations in a sequence of high-frame-rate images and computing local radial symmetry (radiality) around each pixel to estimate the presence of fluorophores with high precision. In this study, SRRF was employed to visualize euchromatin and heterochromatin structures in Solanum lycopersicum root apex cells. Imaging was performed on a Nikon Ti-E inverted microscope equipped with a 100× Plan Apo λ oil immersion objective (NA 1.45) and an Andor iXon Ultra 897U EMCCD camera. For each sample, 3000–5000 image frames were acquired with an exposure time of 50 ms at ~20 frames per second using Andor Solis software. Post-acquisition, SRRF reconstruction was performed using the open-source **NanoJ-SRRF plugin** in **ImageJ/Fiji.** Optimized SRRF parameters included a ring radius of 0.5, radiality magnification of 5, and 6 axes per ring. These parameters allowed pixel interpolation at a 5×5 resolution, effectively increasing the spatial resolution beyond the diffraction limit (~50–70 nm in-plane resolution). The SRRF-processed images revealed high-resolution chromatin domains and nuclear substructures, which were not resolvable using standard confocal microscopy. Quantitative analysis of these SRRF images such as domain number, size, eccentricity, and intensity distribution was conducted using ImageJ’s particle analysis and custom MATLAB scripts.

**Image Analysis**

- **Fluorescence Intensity Quantification**: Region-of-interest (ROI) measurements were conducted using Fiji/ImageJ. Box plots were generated from average pixel intensities (n = 8–10 roots per condition).
- **Line Profile Analysis**: Line scans were drawn across mitochondria, nuclei, or ROS regions to generate intensity distribution curves.
- **Nuclear Metrics**: Nuclear area, DAPI intensity, and chromatin condensation were quantified using automated thresholding and particle analysis.
- **Colocalization Analysis**: Pearson correlation coefficient (PCC), Manders M1 and M2 coefficients between nuclear (DAPI) and NRF2 or mTOR channels were calculated using the Coloc2 plugin in ImageJ.
- **Lysosomal Analysis**: LTR-positive vesicle number and total fluorescence area were quantified using threshold-based segmentation.
- **Chromatin Domain Analysis**: H3K4me3 domain number, area, and eccentricity were calculated using particle analysis on SRRF-processed images.

**Statistical Analysis**

Statistical analyses were performed using SPSS Statistics software to evaluate the significance of differences between treatment groups. Each experimental condition included a sample size of 8–10 root replicates. Data were analyzed using paired two-tailed Student’s *t*-tests to assess statistical significance. A threshold of *p* < 0.05 was used to denote significance, with *p*-values represented as follows: *p* < 0.05. Statistically significant differences were marked accordingly. This approach ensures rigorous assessment of experimental outcomes and enhances the reliability and interpretability of the results. Graphs were generated using GraphPad Prism or OriginPro software.

**Supplementary figures**

1. **Supplementary Figure 1.** Copper reduces tomato seedling growth. Root and shoot length decline with rising Cu doses. Graphs show reduced chlorophyll and root length. *n = 10; mean ± SEM.*
2. **Supplementary Figure 2.** mTOR relocalizes from cytoplasm to nucleus under Cu stress. Co-localization with DAPI increases at moderate doses, then declines. *Scale: 50 µm.*
3. **Supplementary Figure 3.** LTR staining shows dose-dependent lysosome accumulation in root tips, suggesting autophagosome formation near damaged mitochondria.
4. **Supplementary Figure 4.** Hoechst confirms nuclear shrinkage and chromatin condensation. Nuclear area and circularity drop with Cu stress. COM shifts support chromatin marginalization.

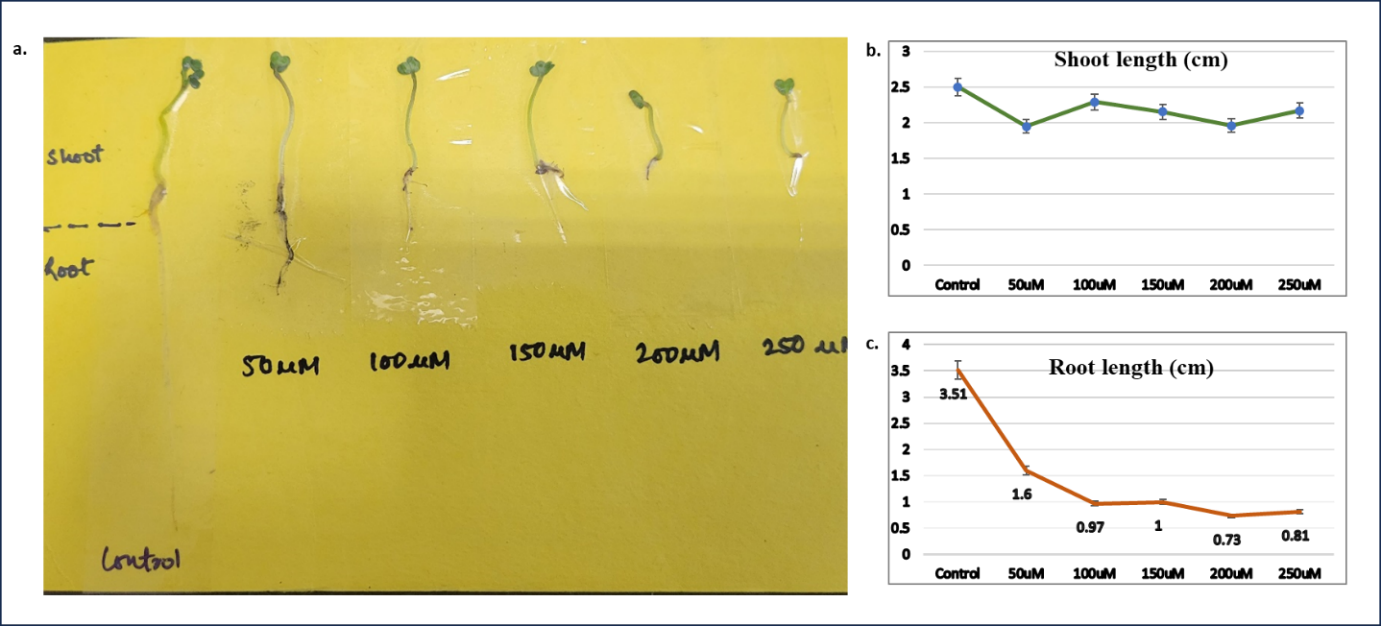

**Supplementary Figure 1.** Copper exposure reduces tomato seedling growth. Images show dose-dependent inhibition of root and shoot elongation. Graphs quantify chlorophyll content and root length, both declining with increasing Cu concentrations. Data represent mean ± SEM (n = 10).

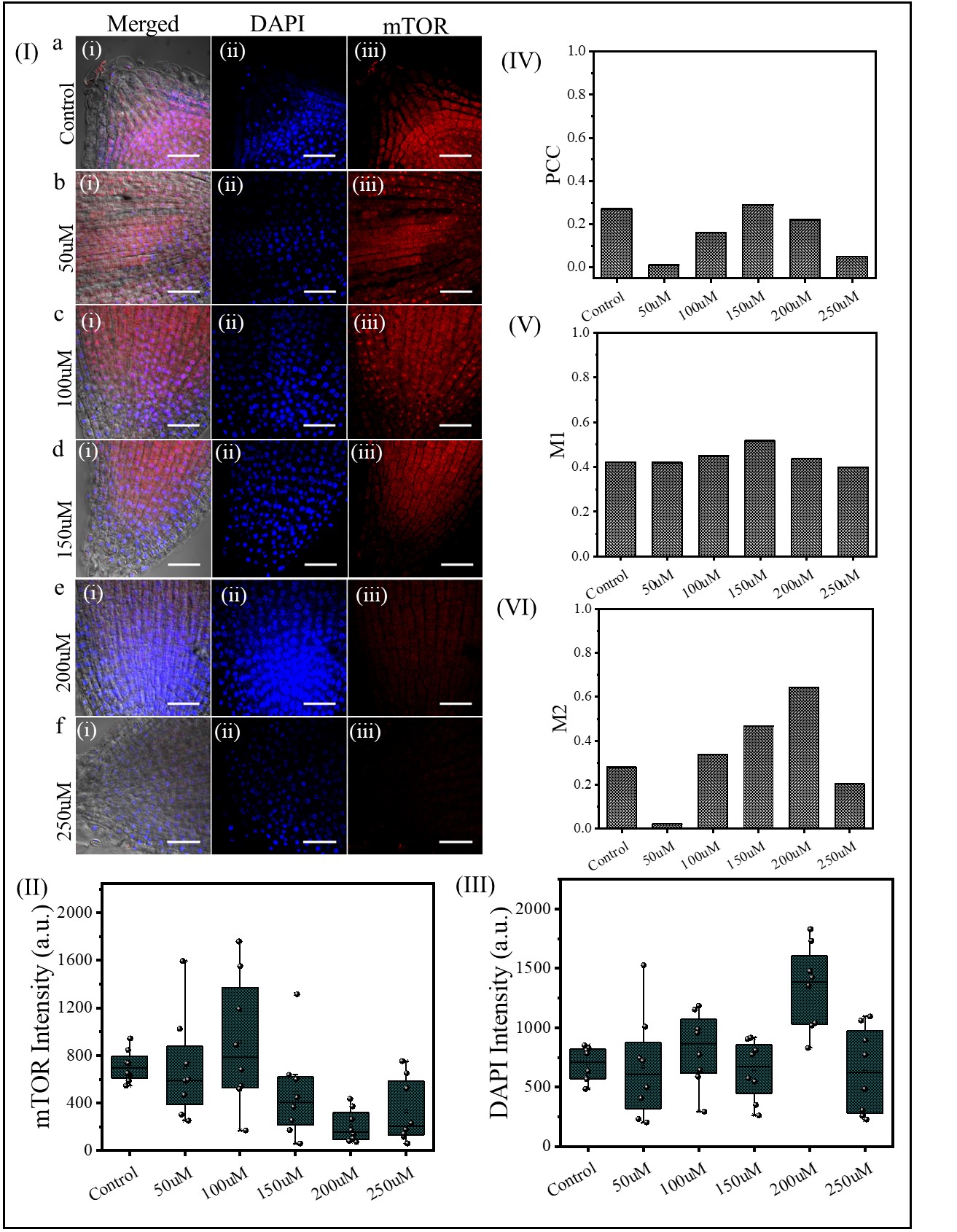

**Supplementary Figure 2.** Confocal analysis of mTOR (red) and DAPI (blue) co-localization in tomato root apex cells under copper stress. (I) Representative images show progressive nuclear condensation and altered mTOR distribution from cytoplasm to nucleus. (II–III) Box plots show changes in mTOR and DAPI fluorescence intensities with increasing Cu concentrations. (IV–VI) Co-localization metrics (PCC, M1, M2) indicate enhanced nuclear association of mTOR under moderate Cu stress, followed by reduction at higher doses. Scale bars: 50 µm.

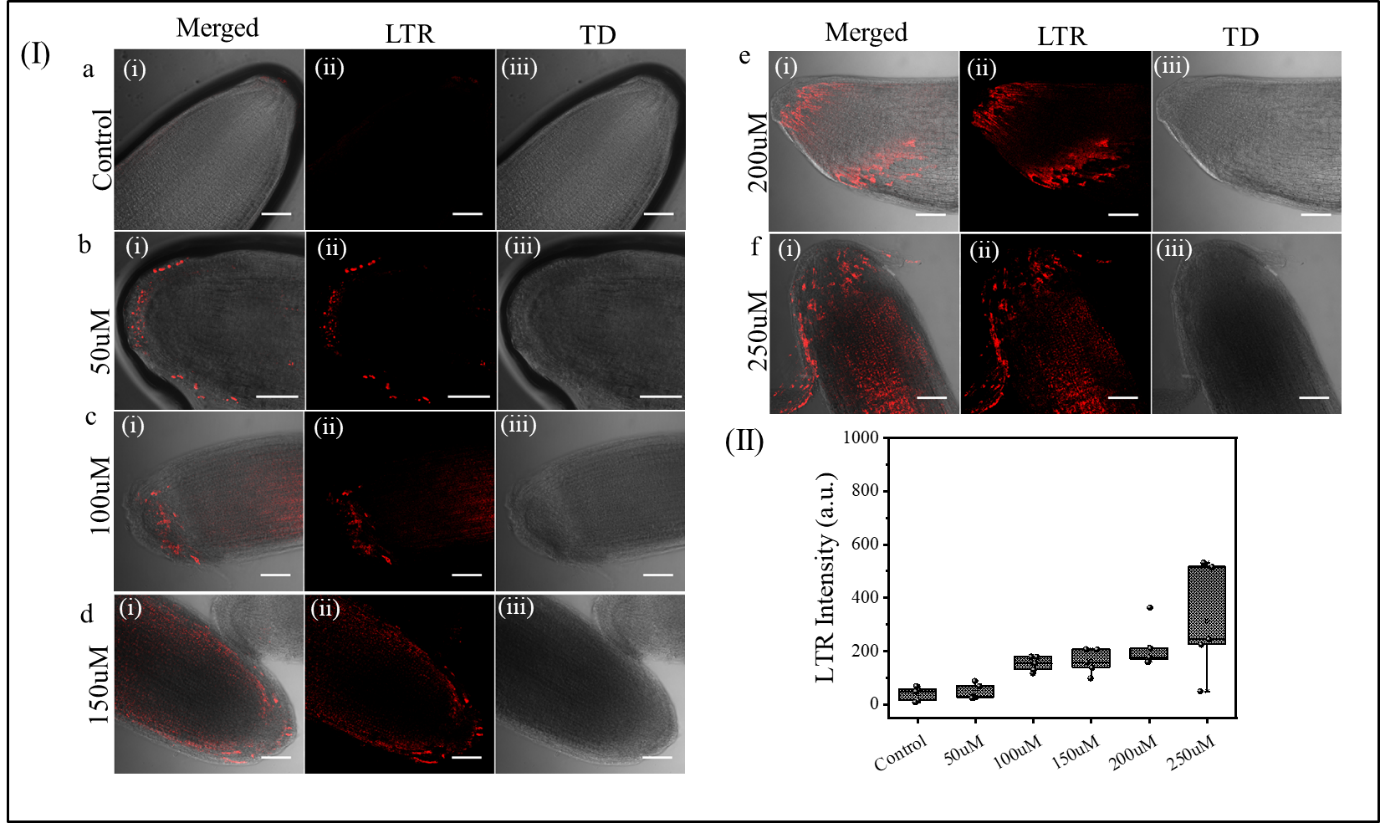

**Supplementary Figure 3.** Autophagosome formation under copper stress in tomato root apex cells visualized using LysoTracker Red (LTR). (I) Confocal images show minimal lysosomal signal in control, with progressive lysosome accumulation and clustering (red) from 50 µM to 250 µM Cu treatment. LTR signal intensifies near the root meristem and elongation zones, especially adjacent to damaged mitochondria. (II) Quantification of LTR fluorescence intensity shows a significant, dose-dependent increase in lysosomal activity under stress. Scale bars: 50 µm. Data represent mean ± SEM, n = 10 root tips per group.

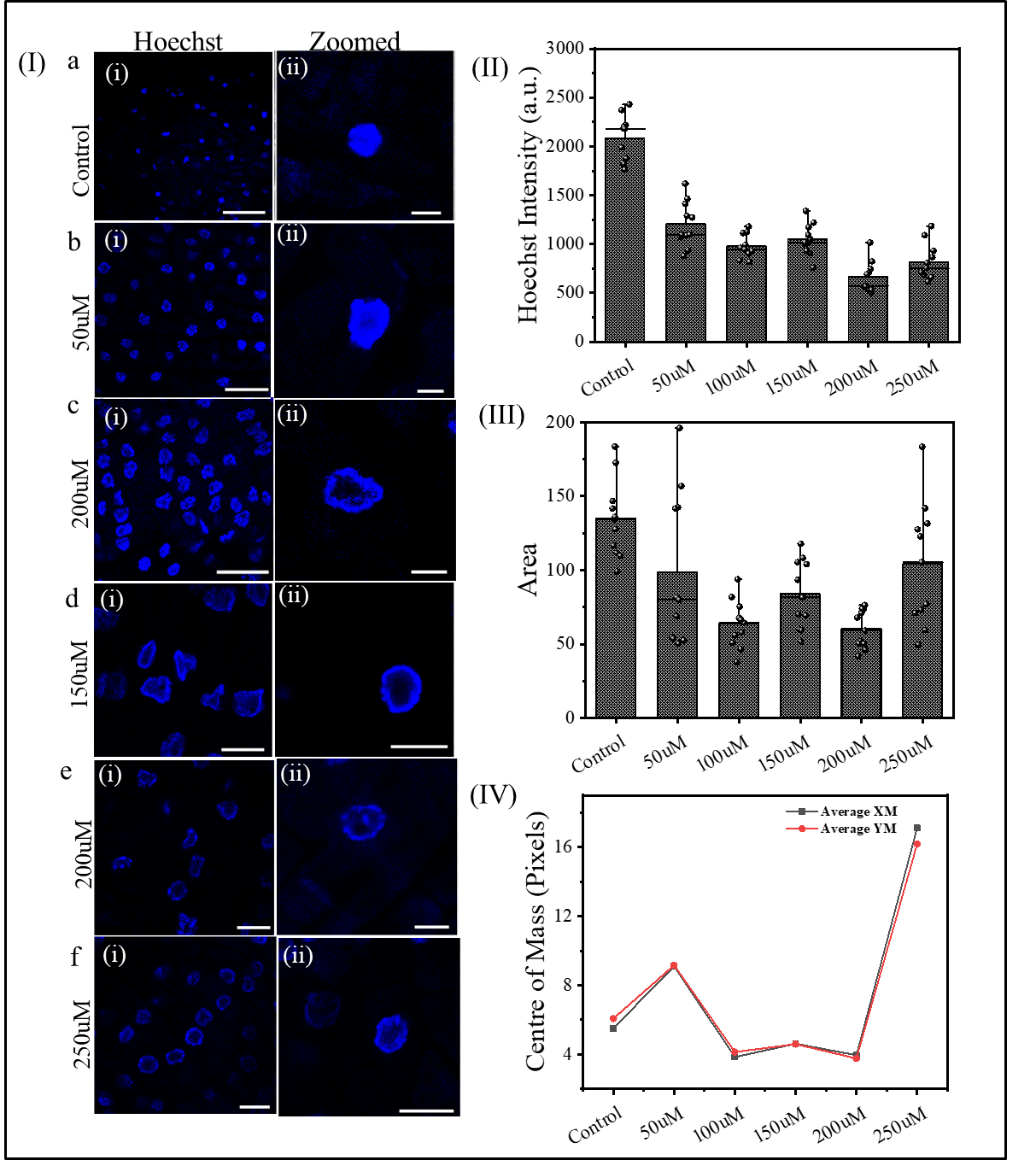

**Supplementary Figure 4.** Validation of nuclear condensation and chromatin reorganization using Hoechst staining under copper stress in tomato root apex cells. (I) Representative images show uniform, round nuclei in control (top), which become increasingly irregular, shrunken, and peripherally condensed with rising Cu concentrations (50–250 µM). (II–IV) Quantitative analysis indicates a decline in nuclear area and circularity, along with shifts in center of mass (COM) coordinates, consistent with nuclear deformation and chromatin marginalization. Scale bars: 50 µm (i), 2 µm (ii). Data are mean ± SEM, n = 10 nuclei per treatment.

.
